# Deciphering the Genomes of Motility-Deficient Mutants of *Vibrio alginolyticus* 138-2

**DOI:** 10.1101/2023.09.26.559574

**Authors:** Kazuma Uesaka, Keita Inaba, Noriko Nishioka, Seiji Kojima, Michio Homma, Kunio Ihara

## Abstract

The motility of *Vibrio* species plays a pivotal role in their survival and adaptation to diverse environments, and is intricately associated with pathogenicity in both humans and aquatic animals. Numerous mutant strains of *Vibrio alginolyticus* have been generated using UV or EMS mutagenesis to probe flagellar motility using molecular genetic approaches. Identifying these mutations promises to yield valuable insights into motility at the protein structural physiology level. In this study, we conducted an exhaustive analysis of the genomic structures of laboratory *Vibrio* strains that served as reference specimens. This assemblage notably encompassed the progenitor strain *V. alginolyticus* 138-2, alongside two strains displaying deficiencies in the lateral flagella (VIO5 and YM4) and one evincing a deficiency in the polar flagella (YM19). Subsequently, we meticulously ascertained the specific mutation sites within the 18 motility-deficient strains whose parents could be traced back to VIO5 or YM4. The identified mutations were conclusively localized within well-established flagella-related genes, providing a coherent rationale for the phenotypic traits exhibited by each motility-deficient mutant. In addition to these motility-related mutations, an average of 20 additional mutations was detected in each mutant strain. Genomic analysis of many mutants is a highly effective tool for comprehensively identifying genes related to specific phenotypes using forward genetics.

**IMPORTANCE:** Bacteria of the *Vibrio* species rely on their motility for survival, adaptation, and pathogenicity. Identifying specific changes in the genome of non-motile mutants can help understand the genes that regulate flagella and motility in these bacteria. Herin, we sequenced the genome of *Vibrio alginolyticus* 138-2 strain and other motility defective strains and identified most of the mutations to be within genes encoding flagella. Additionally, we also provide a coherent rationale for the motility deficiency in these mutants. We believe that this study could provide a framework for genome sequencing-based methods for studying molecular functions in bacteria.

## INTRODUCTION

In biology, genetic approaches assume paramount significance as a method for selecting mutants with specific phenotypic alterations from a vast pool of mutants, concurrently facilitating the identification of the genes responsible for these phenotypes. For instance, the motility apparatus of the marine bacterium *Vibrio alginolyticus* encompasses two flagellar systems: a single polar flagellum expressed constitutively and multiple lateral flagella induced by shifts in the external environment (1-3). To comprehensively understand the distinct functions of these two flagellar types in motility, we generated mutant strains with deficiencies in either the polar or lateral flagella using EMS and/or UV mutagenesis (4, 5). It is noteworthy that subjecting mutants with polar flagella only to EMS treatment yielded a multitude of motility-deficient strains, furthering our understanding of the molecular structure and function of the polar flagella (6).

Genes governing flagellar traits in *V. alginolyticus* and its closely related counterpart, *V. parahaemolyticus*, are clustered within several regions of the genome (7, 8). Nevertheless, it is pertinent to mention that other genes associated with flagella formation have been reported outside of these demarcated regions (9, 10). A comprehensive understanding requires a whole-genome analysis of mutants encompassing genes with broader functionality, including those involved in the assembly and positioning of flagellar structures.

In microbial genome analysis, it has been established that short-read sequencing alone permits simultaneous analysis of approximately 300 strains at an approximate cost of USD 3 per strain (11). The number of contigs resulting from the assembly may vary depending on the target microbial species; however, in the case of *Vibrio*, assembly of N_90_ with more than 100 kb, typically yielding fewer than 100 contigs, is achievable (12-15). Furthermore, the integration of long-read sequencing enables the relatively straightforward acquisition of complete genomes (16-18). Nevertheless, it should be noted that using long-read technologies, such as PacBio or Oxford NanoPore, entails a substantial cost, often amounting to several hundred dollars per strain.

In this study, we devised a method that exclusively utilizes short-read sequencing for genome assembly. This approach was used to elucidate the complete genome structure of *V. alginolyticus* strain 138-2, a wild-type strain featuring a dual flagellar system, two lateral flagellar-deficient strains (VIO5 and YM4), and a polar flagellar-deficient strain (YM19). Additionally, for the 17 mutant strains derived from the aforementioned flagellar-deficient strains, mapping to the genome of the fully characterized parental strain enabled a comprehensive analysis encompassing all mutation types (Single Nucleotide Variations [SNVs], small insertions/deletions [indels], short tandem Repeat Number Variations [RNVs], and large structural variations [LSVs]) and quantities resulting from EMS or UV mutagenesis. The insights gained from this analysis regarding the types and quantities of mutations induced by EMS or UV treatment provide valuable guidance for future mutant generation. Moreover, this analysis focused on mutants linked to flagellar expression systems, particularly those exhibiting chemotaxis deficiencies (Che-type mutants), leading to the accumulation of mutations in genes integral to regulating flagellar rotation. Combining random mutagenesis and expression-based selection provides crucial insights into the efficiency of acquiring target gene mutations.

## RESULTS AND DISCUSSION

### Genome structures of *V. alginolyticus* strain 138-2 and derived mutant strains

We determined the complete genome structure of *V. alginolyticus* strain 138-2 and three mutant strains: VIO5, YM4, and YM19. These strains have been widely used for the functional analysis of polar flagella of the genus *Vibrio* (4). Similar to the genomes reported for other *Vibrio* spp., the genomic DNA consists of two circular chromosomes and does not harbor any plasmids (19). The genome size was 5,185,395 bp for strains 138-2 and VIO5 and 5,185,324 bp for strains YM4 and YM19. DFAST annotation predicted 4,601 protein-coding sequences (CDSs) for strain VIO5, 4,602 for 138-2, and 4,603 for YM4 and YM19. Thirty-seven rRNA genes (twelve 16S rRNA, twelve 23S rRNA, and thirteen 5S rRNA) and one hundred and sixteen tRNA genes were assigned to all four strains (Supplementary Table 1).

### Variation sites in *V. alginolyticus* strains VIO5, YM4, and YM19

*V. alginolyticus* strain VIO5 is a lateral flagellar-deficient mutant that arises from EMS mutagenesis of VIK4 (rifampicin-resistant), which results from a spontaneous mutation in the parent strain 138-2. The rifampicin-resistant phenotype of VIO5 can be attributed to a mutation in chromosome I (position 3,206,619) of the VIO5 genome (Table 1). This mutation leads to a Q513L amino acid substitution in the RNA polymerase beta subunit, which has been reported as a causal SNP of rifampicin resistance in *E. coli* (20). Another mutation on chromosome II (position 1,534,464) introduced a base substitution, causing the 64th tryptophan residue of the MotY2 protein to be replaced by a stop codon. Given that this is the sole mutation observed in chromosome II of VIO5, and considering that all lateral flagellar genes are present on chromosome II, it is highly plausible that this mutation in the *motY2* gene is responsible for lateral flagellar deficiency in the VIO5 strain. Intriguingly, a single mutation in a structural gene can result in the complete loss of flagellar gene expression. The *motY2* gene was one of the earliest members to be expressed in the lateral flagellar expression hierarchy and was positioned at the head of the operon (Fig. 2 B). Therefore, a mutation leading to a premature stop codon in the *motY2* might significantly impact the translation of downstream *lafK* genes, a σ^54^-dependent regulator required for the expression of Class 2 genes of lateral flagella (8).

**Table 1.** Variation sites in the three mutant strains compared to the wild-type strain 138-2.

| position | 138-2 | VIO5 | YM4 | YM19 | mutation | annotation | description |
| --- | --- | --- | --- | --- | --- | --- | --- |
| **Chr. I** |  |  |  |  |  |  |  |
| 135,143 | C | T | C | C | D100N (GAT→AAT) | hslO | 33 kDa chaperonin |
| 165,966 | T | T | A | A | D328V (GAT→GTT) | Vag1382_01380 | bifunctional GTP diphosphokinase<br>guanosine-3',5'-bis(diphosphate) 3'-diphosphatase |
| 213,963 | G | G | C | G | Q180E (CAA→GAA) | Vag1382_01870 | 3-deoxy-D-manno-octulosonic acid transferase |
| 415,183 |  |  | Δ11 bp | Δ11 bp | Y397-FS | tolC | outer membrane channel protein |
| 485,114 | A | C | A | A | I137S (ATC→AGC) | Vag1382_04350 | glutamate synthase |
| 582,644 | A | A | C | C | M256L (ATG→CTG) | phoR | PAS domain-containing sensor histidine kinase |
| 592,588 | (TCAT)<sub>2</sub> | (TCAT)<sub>2</sub> | (TCAT)<sub>1</sub> | (TCAT)<sub>1</sub> | I240-FS | pstB1 | phosphate import ATP-binding protein PstB 1 |
| 1,174,956 | G | G | T | T | A16E (GCA→GAA) | Vag1382_10490 | DNA-binding protein |
| 1,787,856 | (ACCAA)<sub>5</sub> | (ACCAA)<sub>5</sub> | (ACCAA)<sub>6</sub> | (ACCAA)<sub>6</sub> | T451-FS | Vag1382_16050 | hypothetical protein |
| 1,885,712 | (TGTTTT)<sub>2</sub> | (TGTTTT)<sub>2</sub> | (TGTTTT)<sub>1</sub> | (TGTTTT)<sub>1</sub> | intergenic (-206/-323) | ihfA<br>Vag1382_16940 | integration host factor subunit alpha<br>membrane protein |
| 2,065,000 |  |  | Δ57 bp | Δ57 bp | Δ19aa | Vag1382_18450 | membrane protein |
| 2,289,413 | (GCTCTG)<sub>9</sub> | (GCTCTG)<sub>9</sub> | (GCTCTG)<sub>10</sub> | (GCTCTG)<sub>10</sub> | PE repeat 9→10 | Vag1382_20480 | chemotaxis protein CheW |
| 2,299,407 | G | G | G | A | Q295* (CAG→TAG) | flhA | flagellar biosynthesis protein FlhA |
| 2,585,439 | T | T | G | G | Q73H (CAA→CAC) | Vag1382_23340 | hypothetical protein |
| 2,701,438 |  |  | +TA | +TA | I198FS | sfsA | sugar fermentation stimulation protein |
| 3,204,405 | C | C | T | T | S1251N (AGC→AAC) | rpoB | DNA-directed RNA polymerase subunit beta |
| 3,206,619 | T | A | T | T | Q513L (CAG→CTG) | rpoB | DNA-directed RNA polymerase subunit beta |
| **Chr.II** |  |  |  |  |  |  |  |
| 255,362 | G | G | A | G | G53S (GGC→AGC) | flgI2 | flagellar P-ring protein 2 FlgI |
| 513,122 | (AAAAAT)<sub>3</sub> | (AAAAAT)<sub>3</sub> | (AAAAAT)<sub>2</sub> | (AAAAAT)<sub>2</sub> | intergenic (+160/-50) | Vag1382_35310<br>Vag1382_35320 | MFS transporter/methyl-accepting chemotaxis protein |
| 1,534,464 | C | T | C | C | W64* (TGG→TGA) | Vag1382_44100 | flagellar protein MotY2 |

*V. alginolyticus* strain YM4, a lateral flagellar-deficient mutant, and strain YM19, a polar flagellar-deficient mutant, were generated through UV mutagenesis of strain 138-2 (YM19 was further obtained by the spontaneous restoration of the lateral flagellar assembly from a UV-mutagenized strain YM17, which is both a polar and lateral flagellar-deficient mutant). Thirteen mutations were common between YM4 and YM19, suggesting that these mutations accumulated during the early stages of UV mutagenesis (Table 1). Since YM4 and YM19 exhibit different phenotypes for the two types of flagella, a mutation on chromosome II (position 255,362), exclusive to YM4, leading to a change in the 53rd glycine to serine in the lateral flagellar P-ring protein FlgI, is the likely cause of the YM4 lateral flagellar-deficient phenotype. Similarly, a mutation on chromosome I (positions 2,299,407), found only in YM19, mutated the 295th glutamine of the polar flagellar export apparatus protein FlhA to a termination codon, presumably resulting in incomplete FlhA and the polar flagellar-deficient phenotype of YM19. The underlying reason for YM4’s lateral flagellar deficiency is a structural gene mutation (a point mutation that converts glycine to serine). Considering that this gene belongs to one of the operons within class 2 of the lateral flagellar expression hierarchy (8), it is suggested that lateral flagellar expression undergoes subtle regulatory control.

The VIO5 strain generated by EMS mutagenesis had only four SNVs with no insertion or deletion mutations, but the YM4 and YM19 strains created by UV mutagenesis had two deletion mutations and five repeat number variation mutations, in addition to seven and six SNVs for YM4 and YM19, respectively (Table 1).

### Variation sites in other motility-deficient mutant strains

*Vibrio* strains NMB136, NMB155, and KK148 were generated from strain VIO5, whereas strains NMB75, NMB82, NMB88, NMB93, NMB95, NMB98, NMB99, NMB102, NMB103, NMB105, NMB106, NMB111, and NMB116 were generated from strain YM4. All 16 strains were generated through EMS mutagenesis and screened as mutants that could not form a swimming ring on a soft agar plate; motility was observed under dark-field microscopy (21). Supplementary Table S2 summarizes the detected mutation sites in the genome of each strain compared with the 138-2 strain genome. The mutations were categorized into three types: single nucleotide variations (SNVs), short insertions/deletions (indels), and short tandem repeat number variations (RNVs). Considering these variations, the number of SNVs, indels, and RNVs detected in each strain was used to create a pedigree for the strain (Figure 1). Because NMB136, NMB155, and KK148 were generated from the VIO5 strain at different times, these three strains carried completely independent mutations. Conversely, the 14 NMB strains generated from YM4 almost simultaneously carried nine common mutations (5 SNVs, 2 indels, and 2 RNVs) in addition to various unique mutations ranging from 2 to 54. All variations from the 17 strains created by EMS mutagenesis and the 3 strains created by UV mutagenesis are summarized in Table 2. The analysis showed that UV mutagenesis produced more indels and RNVs than SNVs. In contrast, most EMS mutations (227/237) were SNVs. Among the SNVs with EMS mutations, 209 were found within protein-coding genes, 17 in intergenic regions, and 1 in tRNA. Among the 209 SNVs in the protein-coding genes, 159 were non-synonymous substitutions, and 59 were synonymous substitutions, resulting in a non-synonymous to synonymous ratio of 2.69, which reflected the effects of non-selective random mutations. Additionally, among the SNVs, 207 showed G:C to A:T transitions, a prominent characteristic of EMS mutagenesis (Table 3). Considering the individual strains analyzed, the mutations they carried were counted independently, resulting in an average of 20.6 ± 12.7 mutations (mean ± standard deviation).

**FIG 1.**
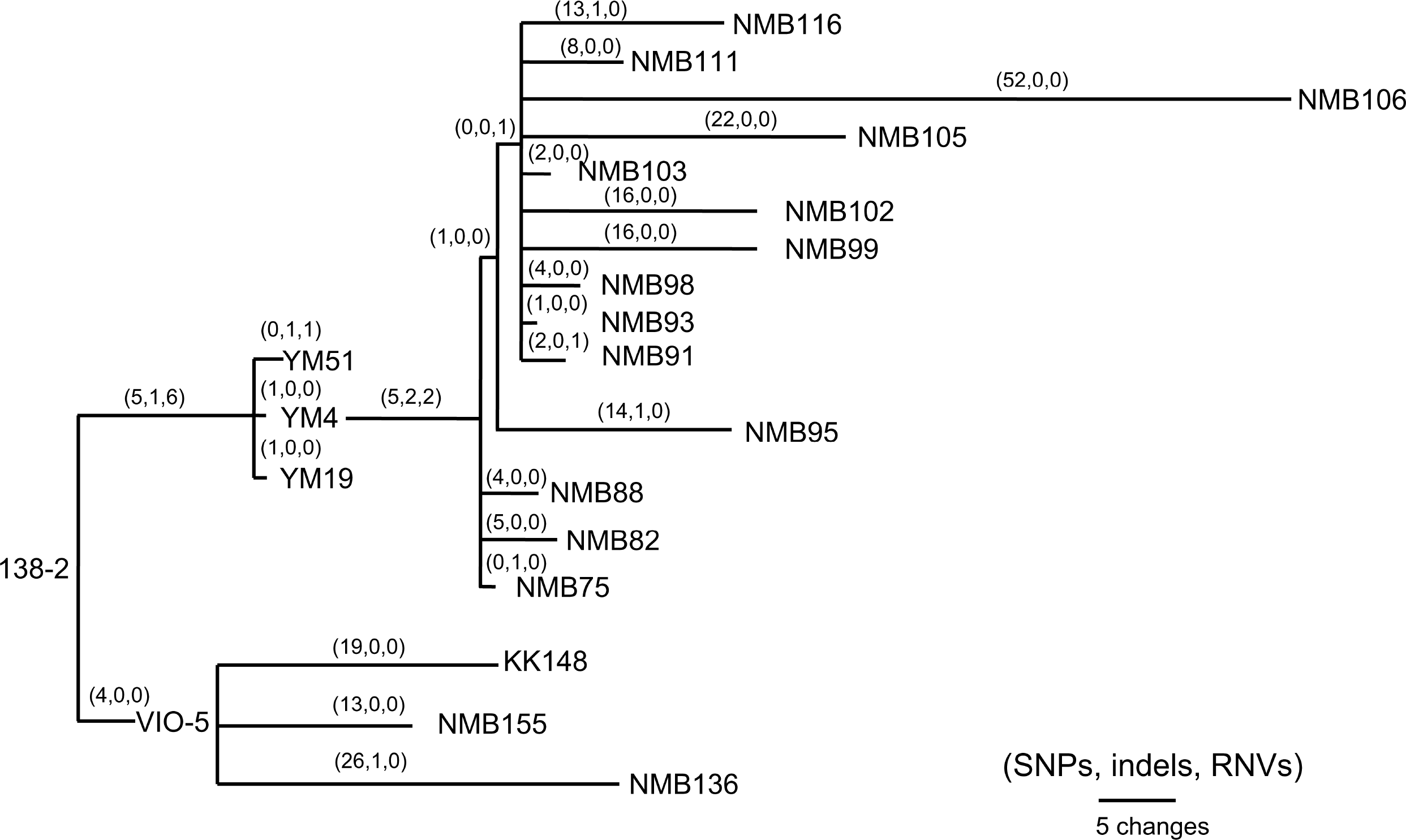
Pedigree tree based on the SNVs, indels, and RNVs among *V. alginolyticus* mutant strains. Illustration of the phylogeny of the mutant strains, assuming the mutations (SNVs, indels, and RNVs) to be equidistant. The three numbers in parentheses drawn above the branches represent SNVs, indels, and RNVs. One mutation was presumed to have occurred later in the YM4 strain than when it was used to create the NMB strain.

**Table 2.**
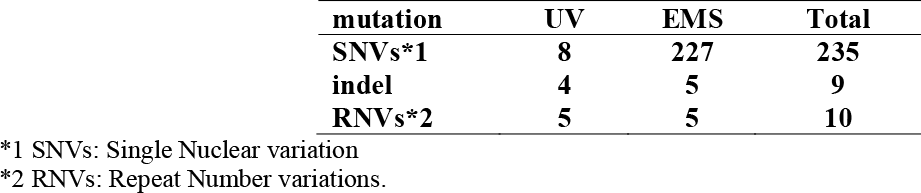
Frequency of different mutations found in EMS or UV mutagenesis.

**Table 3.**
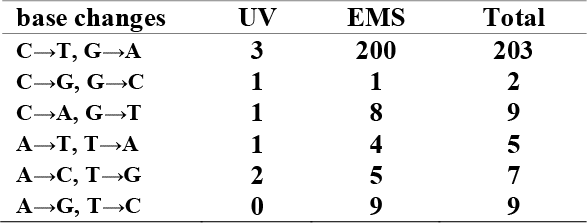
Frequency of base changes in SNVs found in all EMS or UV mutants.

| **base changes** | **UV** | **EMS** | **Total** |
| --- | --- | --- | --- |
| **C→T, G→A** | **3** | **200** | **203** |
| **C→G, G→C** | **1** | **1** | **2** |
| **C→A, G→T** | **1** | **8** | **9** |
| **A→T, T→A** | **1** | **4** | **5** |
| **A→C, T→G** | **2** | **5** | **7** |
| **A→G, T→C** | **0** | **9** | **9** |

### Putatively responsible variations for motility-deficient mutant strains

The motility-deficient mutants analyzed in this study can be classified into three types: those with no or incomplete flagella (Fla^−^ type), those with chemotaxis problems (Che^−^ type), and those with an increased number of polar flagella (Pof^m^ type). Another type, the Mot^−^ type, which has abnormalities in the rotation apparatus, was absent in the analyzed mutants. These three types of motility-deficient mutants had 10–30 variation sites, but all had mutations in a known flagella-related gene (Fig. 2A and B). The Fla^−^ type mutants, NMB103 and NMB116, had mutations in the *flgL* gene, leading to the observation of only the hook structure without visible flagellar filaments, which explains their flagella-deficient phenotype. Che^−^ type mutants included NMB75, NMB82, NMB88, NMB91, NMB93, NMB95, NMB98, NMB99, NMB102, NMB105, NMB106, NMB111, and NMB136. NMB82 and NMB105 harbored mutations in the *cheA* gene, NMB91 and NMB98 harbored mutations in the *zomB* gene, NMB93 and NMB136 harbored mutations in the *cheY* gene, and NMB88, NMB95, NMB99, NMB102, and NMB106 harbored mutations in the *fliM* gene. These genes are part of the gene cluster responsible for the chemotactic response in *Vibrio*, especially the change in flagellar rotation (see Supplementary Fig. S2). Each strain exhibits various degrees of fixed rotation in its expression system. Among the Che^−^ type mutants, the *fliM* gene, identified as the causal gene for many mutations, appears to be involved in flagellar rotation control and the regulation of flagellar numbers (22). This suggests versatile roles of the *fliM* gene in governing flagellar expression. The NMB75 strain harbored a mutation in the *cheR* gene, resulting in a leaky phenotype due to partial signal transmission by CheA, partially affected by the deficiency of CheR activity in methylating chemoreceptors. NMB111 showed weakly reduced swimming ability due to mutations in the *flhG* gene, which regulates flagellar number, and flagella were rarely observed with the FlhG(D171A) mutation (23). The reduced swimming ability of NMB111 may be due to the reduced flagellar number caused by the FlhG(D171N) mutation. Thus, the NMB111 strain may be included in the Fla^−^ type mutants.

**FIG 2.**
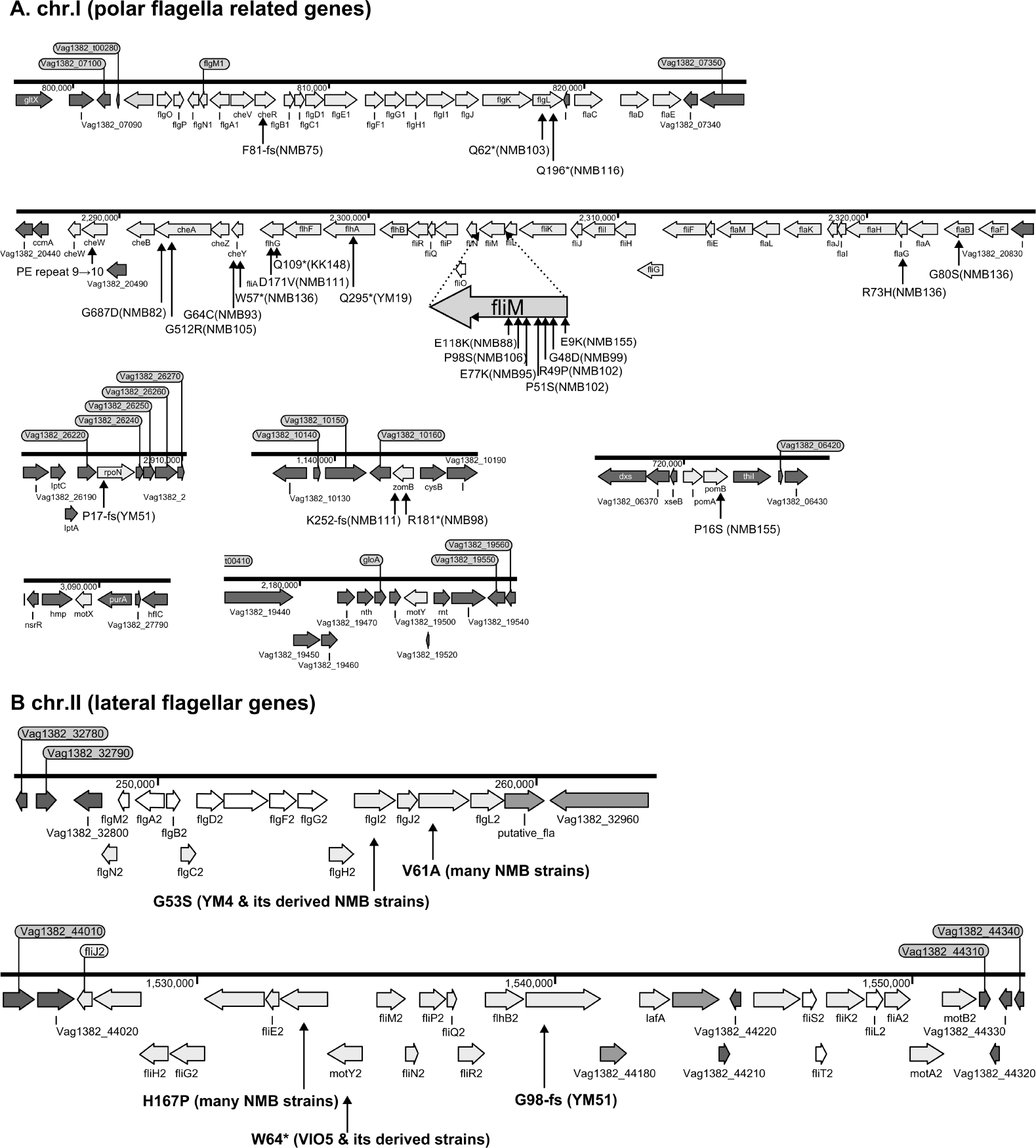
Known flagella-related genes in chromosome I (A) and chromosome II (B) and the detected variations in mutant strains. Chromosome I contains a cluster of genes related to the polar flagellum and chemotaxis signal transduction (A), and chromosome II contains a cluster of genes related to the lateral flagella (B). Chromosome I consists of seven regions: two regions consisting of large clusters and five regions consisting of one or two genes; chemo-signal transduction genes are found in one of the two large clusters. In addition to this, chemoreceptors are present scattered throughout the genome. Chromosome II consists of two large clusters in two regions. Known genes are listed by their customary names, and the paralogs of the lateral hair genes on chromosome II are distinguished by the four letters, followed by the number 2. The arrow extended below the gene indicates the location of the mutation that occurred in that gene. A mutation is an amino acid change found in the protein, and the name of the strain in which the mutation was found is provided in parentheses.

Pof^m^ type mutants included the KK148 and NMB155 strains. The KK148 strain harbored a mutation (Q109*) in the *flhG* gene, as previously reported (24), whereas the NMB155 strain harbored a mutation (E9K) in the *fliM* gene, as previously reported (22). In KK148, a defective *flhG* gene product has been shown to form multiple polar flagella (24). Similarly, it has been shown that the FliM(E9K) mutation changes flagellar numbers (22). Based on the evidence, the fact that the *fliM* gene, which has been implicated in chemotaxis, also plays a significant role in regulating flagellar number in the Che^−^ type mutants is highly intriguing. YM51 was isolated as a motility-deficient mutant of YM5 and exhibits a Fla^−^ type expression system. Interestingly, unlike YM4, the reason for YM51’s lateral flagellar suppression was found to be a frame-shift mutation in the *flhA* gene (Fig. 2B) in a region where seven consecutive G bases were found to have increased by one, causing truncation of the FlhA protein to approximately 100 amino acids. In *Paenibacillus glucanolyticus*, it was recently reported that swarming was suppressed by reversible hotspots that reduced the number of eight consecutive A bases to seven and that the strain could easily revert to a swarm-competent state (25). In lateral flagella-deficient mutants of *Vibrio*, there appear to be two types of lateral flagella-deficient Vibrio mutants: a strong phenotype with almost irreversible mutations, such as VIO5 and YM4, and a leaky phenotype that is relatively easy to revert, such as YM51. A causal mutation in YM5’s lateral flagellar-deficient phenotype may be the same as in strain YM51. The YM51 strain, similar to NMB103 and NMB116, showed only a hook structure without visible flagellar filaments, indicating that the assembly of the flagellar filament was impaired (26). However, no mutations were found in flagellar structure genes, including the *flgL* gene, but a mutation was detected in the *rpoN* gene, which plays an important role in polar flagella formation (27). Although the YM14 strain used for cloning the *rpoN* gene was not included in the current genome analysis, it is highly plausible that YM51 has a mutation similar to that of YM14.

Through genomic analysis of 17 different mutants selected by a combination of EMS mutagenesis and screening for motility-deficient phenotypes, we found that these strains contained 3–75 different gene mutations, in addition to one or two genes presumed to be responsible for the phenotype. The results of this genome analysis demonstrated that it is possible to statistically identify responsible genes by analyzing a certain number of mutant strains, as it is not possible to narrow down the responsible genes by analyzing individual mutants alone. Furthermore, the analysis of only 17 strains revealed the extraction of 7 genes involved in the control system of flagella rotation (Supplementary Fig. S3), suggesting that increasing the number of analyzed mutants can comprehensively extract genes in the entire system. Although Genome Wide Association Studies (GWAS) have rarely been conducted for genetically manipulable microorganisms, if a sufficient number of mutant strains can be created for microorganisms handled in the laboratory, GWAS analysis can effectively detect components of various biological systems.

## MATERIALS AND METHODS

### Bacterial cultivation and genomic DNA isolation

All *Vibrio alginolyticus* strains used in this study are summarized in Supplementary Table S3. Strain VIO5 is a lateral flagella-deficient mutant (Pof^+^, Laf^−^) generated by EMS mutagenesis of VIK4, a spontaneous rifampicin-resistant strain obtained from the wild-type strain 138-2 (Pof^+^, Laf^+^) (5). Strains YM4 and YM5 are lateral flagella-deficient mutants (Pof^+^, Laf^−^) obtained by UV treatment of strain 138-2, whereas strain YM19 is a polar flagellum-deficient mutant (Pof^−^, Laf^+^). This mutant was spontaneously isolated from YM17, a UV-treated YM5-derived strain with impaired polar and lateral flagellar formation (4). Strain YM51 is a motility-deficient mutant of strain YM5, and strains NMB75, 82, 88, 93, 95, 98, 99, 102, 103, 105, 106, 111, and 116 are motility-deficient mutants of strain YM4 (21, 26). Strains NMB136, NMB155, and KK148 are motility-deficient mutants derived from VIO5(24, 28). All *Vibrio* strains were cultivated in VC medium (0.5%(w/v) tryptone, 0.5%(w/v) yeast extract, 0.4%(w/v) K_2_HPO_4_, 3%(w/v) NaCl, and 0.2%(w/v) glucose). Genomic DNA was isolated in the late logarithmic phase using a genomic DNA isolation kit (Promega, WI, USA).

### *De novo* assembly and construction of complete *V. alginolyticus strain* VIO5 genome

*V. alginolyticus* strain VIO5 has been used in many sodium-driven polar flagellar studies (29). In this study, we first determined the complete genome sequence of strain VIO5. Paired-end sequencing reads (2 × 300-bp) were trimmed by quality value, and adapter sequences were removed using Trimmomatics (30). Trimmed reads were *de novo* assembled with SPAdes v.3.1(31). From approximately 150 assembled contigs, we selected 27 contigs larger than 3 kb whose coverage was close to the most frequent value. Primer 3 (32) was used to design forward primers specific to both ends of each contig, except for the repetitive contigs. (Supplementary Table S4). The direction and order of the long contigs were predicted by alignment with the genomic sequences of the most closely related *Vibrio* strain (*Vibrio* sp. EX25; accession numbers NC_13456 and NC_13457) using Mauve software (33). During this operation, the orientation and order of 23 of the 27 contigs were successfully determined. Two contigs formed a circular chromosome (chromosome II) and 21 contigs were linked to form four large assemblies. To determine the linkage order between these four large assemblies and the remaining four contigs that were not aligned in the EX25 genome, PCR experiments were performed using all combinations of primer sets. The amplified PCR fragments were checked for uniformity and size using 0.8% agarose gel electrophoresis and purified using a PCR fragment recovery kit (Promega, WI, USA). The recovered DNA fragments were sequenced using the MiSeq 500PE v.2 kit (Illumina, San Diego, CA, USA). Short-read sequences from each PCR fragment were assembled using Spades v 3.1(31), and the contig with the largest size and highest coverage was adopted as the PCR fragment. For the three regions in which two large contigs appeared after assembly (Supplementary Table S5), it was inferred that the rRNA operons were arranged in tandem. Therefore, we designed primers in both directions in the center of the connecting contig (approximately 200 bp, contig 175 in Table S3) expected to be between these tandemly arranged rRNA operons, and amplified the two rRNA operons independently by PCR. Both the amplified fragments were independently determined and connected using a central contig. Chromosome I of strain VIO5 was completed by manually combining all contigs and PCR fragment sequences. In regions where discrepancies were found between the contig and PCR fragment sequences, the results of the PCR fragment were used preferentially for linkage because the contig ends may contain polymorphisms due to repeat sequences. Finally, sequencing reads were mapped against the full-length genome sequence using BWA (34) to confirm the absence of assembly errors. The Integrative Genomics Viewer (IGV) (35) was used for map visualization.

### Complete genomes of *V. alginolyticus* 138-2, YM4, and YM19, using *V. alginolyticus* VIO5 as a reference strain

Next, we developed a workflow aimed at completing the genome structure of the target *Vibrio* strains in cases where short-read sequencing technology and very closely related reference strains were available; however, long-read sequencing technology was unavailable because of higher sequencing costs or challenges in high molecular weight DNA extraction (Supplementary Fig. S1). This workflow was applied to assemble the complete genome structures of three strains (*V. alginolyticus* strains 138-2, YM4, and YM19) that are closely related to strain VIO5. Paired-end sequencing reads were trimmed based on their quality using fastp v.0.20.0 (36) and used as input data (hereafter, WGS reads). WGS reads were assembled *de novo* using SPAdes v.3.13 (31) to produce error-corrected WGS contigs by two cycles of polishing with Pilon v1. 23 (37). WGS contigs of less than 1 kbp in length or contigs with abnormal coverage were excluded, and the terminal 127-bp of the remaining WGS contigs were trimmed. These contigs were designated long and normal coverage contigs. For the selection method based on the coverage count, the median of the average read counts of the contigs up to the top five in length was set as value C. Contigs whose coverage was more than twice or less than half of value C were removed as abnormal coverage contigs. Bbmap v.37.62, published by JGI(38) was used for coverage calculation. LN contigs with a 16-mer frequency greater than or equal to 2 were hard-masked using Primer3_masker (39). Then, primer3 (40) was used to design outward primers within 1 kb at both ends of each contig. The specificity of the designed primers was checked using FastPCR (41) (Supplementary Table S4). The workflow from short reads to LN contigs and primer design is available on GitHub: script genome_quest (https://github.com/kazumaxneo/genome_quest). Minimap2 (42) was used to align each LN contig with the complete genome sequence of the *V. alginolyticus* VIO5. PCR was performed using primers designed for each LN contig. Multiplexed PCR products were sequenced using MiSeq and individually assembled, as described in the previous section. LN contigs and Locally Assembled PCR fragment (LA) contigs were connected using CAP3 (43.). Two circular chromosomes were also identified. Finally, WGS reads were mapped to the two assembled chromosomal DNA sequences using minimap2 (42), to correct as many errors as possible using the following variation detection tools: breseq (44), GATK HaplotypeCaller v 3.8(45), minimap2 paftool (42), and SV Quest (Uesaka et al., in preparation) (v1.0) (https://github.com/kazumaxneo/SV-Quest).

The complete genome sequences of the four strains have been published in the database under the accession numbers AP022859–AP022866.

### Variation analyses in the mutant strains

Genomic libraries were constructed and sequenced by MiSeq using paired-end sequencing (2 × 300 bp) for 17 strains generated by EMS mutagenesis and one strain, YM51, derived from YM5. Using the genome of *V. alginolyticus* 138-2 as a reference, all variants (SNVs, Indels, RNVs, and LSVs) present in the three strains were extracted using Equation (44) and other tools, as described in the previous section.

### Core gene phylogenetic tree inference

Each mutant genome was created using the gdtools APPLY command from the output of the Breseq variant calling against the 138-2 genome sequence (44). Core genome alignment of the 138-2 sequence and the derived sequence of the mutant strain was performed using Parsnp (https://github.com/marbl/parsnp). Unreliable alignment blocks were excluded based on the Parsnp criteria. A phylogenetic tree was manually constructed based on the number of variations.

## ACKNOWLEDGMENTS

This study utilized mutants developed in a series of studies on the structural and functional analysis of *Vibrio* polar flagella driven by Na^+^ electrochemical potential differences. This research endeavor was initially initiated by Prof. Imae and was subsequently carried forward by two authors, Homma and Kojima. We extend our gratitude to Kawagishi, Okunishi, Maekawa, Kusumoto, and Yorimitsu for constructing the mutants. We would like to thank Editage (www.editage.jp) for English language editing.

## Supplementary Material

Table S1 to S5 (excel)

Fig.S1 to Fig.S3 (pdf)

Fig.S1, S2, and S3 with legends.

## REFERENCES

1. Atsumi T, McCarter L, Imae Y. 1992. Polar and lateral flagellar motors of marine Vibrio are driven by different ion-motive forces. Nature 355:182–4.

2. Atsumi T, Maekawa Y, Yamada T, Kawagishi I, Imae Y, Homma M. 1996. Effect of viscosity on swimming by the lateral and polar flagella of Vibrio alginolyticus. J Bacteriol 178:5024–6.

3. McCarter LL. 2004. Dual flagellar systems enable motility under different circumstances. J Mol Microbiol Biotechnol 7:18–29.

4. Kawagishi I, Maekawa Y, Atsumi T, Homma M, Imae Y. 1995. Isolation of the polar and lateral flagellum-defective mutants in Vibrio alginolyticus and identification of their flagellar driving energy sources. J Bacteriol 177:5158–60.

5. Okunishi I, Kawagishi I, Homma M. 1996. Cloning and characterization of motY, a gene coding for a component of the sodium-driven flagellar motor in Vibrio alginolyticus. J Bacteriol 178:2409–15.

6. Homma M, Nishikino T, Kojima S. 2022. Achievements in bacterial flagellar research with focus on Vibrio species. Microbiol Immunol 66:75–95.

7. Kim YK, McCarter LL. 2000. Analysis of the polar flagellar gene system of Vibrio parahaemolyticus. J Bacteriol 182:3693–704.

8. Stewart BJ, McCarter LL. 2003. Lateral flagellar gene system of Vibrio parahaemolyticus. J Bacteriol 185:4508–18.

9. Yamaichi Y, Bruckner R, Ringgaard S, Moll A, Cameron DE, Briegel A, Jensen GJ, Davis BM, Waldor MK. 2012. A multidomain hub anchors the chromosome segregation and chemotactic machinery to the bacterial pole. Genes Dev 26:2348–60.

10. Brenzinger S, Pecina A, Mrusek D, Mann P, Volse K, Wimmi S, Ruppert U, Becker A, Ringgaard S, Bange G, Thormann KM. 2018. ZomB is essential for flagellar motor reversals in Shewanella putrefaciens and Vibrio parahaemolyticus. Mol Microbiol 109:694–709.

11. Shapland EB, Holmes V, Reeves CD, Sorokin E, Durot M, Platt D, Allen C, Dean J, Serber Z, Newman J, Chandran S. 2015. Low-Cost, High-Throughput Sequencing of DNA Assemblies Using a Highly Multiplexed Nextera Process. ACS Synth Biol 4:860–6.

12. Castillo D, D’Alvise P, Kalatzis PG, Kokkari C, Middelboe M, Gram L, Liu S, Katharios P. 2015. Draft Genome Sequences of Vibrio alginolyticus Strains V1 and V2, Opportunistic Marine Pathogens. Genome Announc 3.

13. Deb S, Badhai J, Das SK. 2020. Draft Genome Sequences of Vibrio alginolyticus Strain S6-61 and Vibrio diabolicus Strain S7-71, Isolated from Corals in the Andaman Sea. Microbiol Resour Announc 9.

14. Meza G, Majrshi H, Saurabh Singh K, Deole R, Tiong HK. 2022. Draft Genome Sequences of the Vibrio parahaemolyticus Strains VHT1 and VHT2, Pasteurization-Resistant Isolates from Environmental Seafood. Microbiol Resour Announc 11:e0079322.

15. LaPorte JP, Spinard EJ, Cavanagh D, Gomez-Chiarri M, Rowley DC, Mekalanos JJ, Mittraparp-Arthorn P, Nelson DR. 2023. Draft Genome Sequence of Vibrio parahaemolyticus PSU5579, Isolated during an Outbreak of Acute Hepatopancreatic Necrosis Disease in Thailand. Microbiol Resour Announc 12:e0087322.

16. Miyamoto M, Motooka D, Gotoh K, Imai T, Yoshitake K, Goto N, Iida T, Yasunaga T, Horii T, Arakawa K, Kasahara M, Nakamura S. 2014. Performance comparison of second- and third-generation sequencers using a bacterial genome with two chromosomes. BMC Genomics 15:699.

17. Chen Z, Erickson DL, Meng J. 2020. Benchmarking hybrid assembly approaches for genomic analyses of bacterial pathogens using Illumina and Oxford Nanopore sequencing. BMC Genomics 21:631.

18. Wick RR, Judd LM, Holt KE. 2023. Assembling the perfect bacterial genome using Oxford Nanopore and Illumina sequencing. PLoS Comput Biol 19:e1010905.

19. Okada K, Iida T, Kita-Tsukamoto K, Honda T. 2005. Vibrios commonly possess two chromosomes. J Bacteriol 187:752–7.

20. Campbell EA, Korzheva N, Mustaev A, Murakami K, Nair S, Goldfarb A, Darst SA. 2001. Structural mechanism for rifampicin inhibition of bacterial rna polymerase. Cell 104:901–12.

21. Homma M, Oota H, Kojima S, Kawagishi I, Imae Y. 1996. Chemotactic responses to an attractant and a repellent by the polar and lateral flagellar systems of Vibrio alginolyticus. Microbiology (Reading) 142 (Pt 10):2777–83.

22. Homma M, Takekawa N, Fujiwara K, Hao Y, Onoue Y, Kojima S. 2022. Formation of multiple flagella caused by a mutation of the flagellar rotor protein FliM in Vibrio alginolyticus. Genes Cells 27:568–578.

23. Ono H, Takashima A, Hirata H, Homma M, Kojima S. 2015. The MinD homolog FlhG regulates the synthesis of the single polar flagellum of Vibrio alginolyticus. Mol Microbiol 98:130–41.

24. Kusumoto A, Kamisaka K, Yakushi T, Terashima H, Shinohara A, Homma M. 2006. Regulation of polar flagellar number by the flhF and flhG genes in Vibrio alginolyticus. J Biochem 139:113–21.

25. Hefetz I, Israeli O, Bilinsky G, Plaschkes I, Hazkani-Covo E, Hayouka Z, Lampert A, Helman Y. 2023. A reversible mutation in a genomic hotspot saves bacterial swarms from extinction. iScience 26:106043.

26. Nishioka N, Furuno M, Kawagishi I, Homma M. 1998. Flagellin-containing membrane vesicles excreted from Vibrio alginolyticus mutants lacking a polar-flagellar filament. J Biochem 123:1169–73.

27. Kawagishi I, Nakada M, Nishioka N, Homma M. 1997. Cloning of a Vibrio alginolyticus rpoN gene that is required for polar flagellar formation. J Bacteriol 179:6851–4.

28. Kojima S, Asai Y, Atsumi T, Kawagishi I, Homma M. 1999. Na+-driven flagellar motor resistant to phenamil, an amiloride analog, caused by mutations in putative channel components. J Mol Biol 285:1537–47.

29. Li N, Kojima S, Homma M. 2011. Sodium-driven motor of the polar flagellum in marine bacteria Vibrio. Genes Cells 16:985–99.

30. Bolger AM, Lohse M, Usadel B. 2014. Trimmomatic: a flexible trimmer for Illumina sequence data. Bioinformatics 30:2114–20.

31. Bankevich A, Nurk S, Antipov D, Gurevich AA, Dvorkin M, Kulikov AS, Lesin VM, Nikolenko SI, Pham S, Prjibelski AD, Pyshkin AV, Sirotkin AV, Vyahhi N, Tesler G, Alekseyev MA, Pevzner PA. 2012. SPAdes: a new genome assembly algorithm and its applications to single-cell sequencing. J Comput Biol 19:455–77.

32. Rozen S, Skaletsky H. 2000. Primer3 on the WWW for general users and for biologist programmers. Methods Mol Biol 132:365–86.

33. Darling AC, Mau B, Blattner FR, Perna NT. 2004. Mauve: multiple alignment of conserved genomic sequence with rearrangements. Genome Res 14:1394–403.

34. Li H, Durbin R. 2009. Fast and accurate short read alignment with Burrows-Wheeler transform. Bioinformatics 25:1754–60.

35. Thorvaldsdóttir H, Robinson JT, Mesirov JP. 2013. Integrative Genomics Viewer (IGV): high-performance genomics data visualization and exploration. Brief Bioinform 14:178–92.

36. Chen S, Zhou Y, Chen Y, Gu J. 2018. fastp: an ultra-fast all-in-one FASTQ preprocessor. Bioinformatics 34:i884–i890.

37. Walker BJ, Abeel T, Shea T, Priest M, Abouelliel A, Sakthikumar S, Cuomo CA, Zeng Q, Wortman J, Young SK, Earl AM. 2014. Pilon: an integrated tool for comprehensive microbial variant detection and genome assembly improvement. PLoS One 9:e112963.

38. JGI. BBMap. https://sourceforge.net/projects/bbmap/. Accessed

39. Kõressaar T, Lepamets M, Kaplinski L, Raime K, Andreson R, Remm M. 2018. Primer3_masker: integrating masking of template sequence with primer design software. Bioinformatics 34:1937–1938.

40. Untergasser A, Cutcutache I, Koressaar T, Ye J, Faircloth BC, Remm M, Rozen SG. 2012. Primer3--new capabilities and interfaces. Nucleic Acids Res 40:e115.

41. Kalendar R, Khassenov B, Ramankulov Y, Samuilova O, Ivanov KI. 2017. FastPCR: An in silico tool for fast primer and probe design and advanced sequence analysis. Genomics 109:312–319.

42. Li H. 2018. Minimap2: pairwise alignment for nucleotide sequences. Bioinformatics 34:3094–3100.

43. Huang X, Madan A. 1999. CAP3: A DNA sequence assembly program. Genome Res 9:868–77.

44. Deatherage DE, Barrick JE. 2014. Identification of mutations in laboratory-evolved microbes from next-generation sequencing data using breseq. Methods Mol Biol 1151:165–88.

45. DePristo MA, Banks E, Poplin R, Garimella KV, Maguire JR, Hartl C, Philippakis AA, del Angel G, Rivas MA, Hanna M, McKenna A, Fennell TJ, Kernytsky AM, Sivachenko AY, Cibulskis K, Gabriel SB, Altshuler D, Daly MJ. 2011. A framework for variation discovery and genotyping using next-generation DNA sequencing data. Nat Genet 43:491–8.

