## Supplementary Figures for "Deciphering the Genomes of Motility-Deficient Mutants of *Vibrio alginolyticus* 138-2"

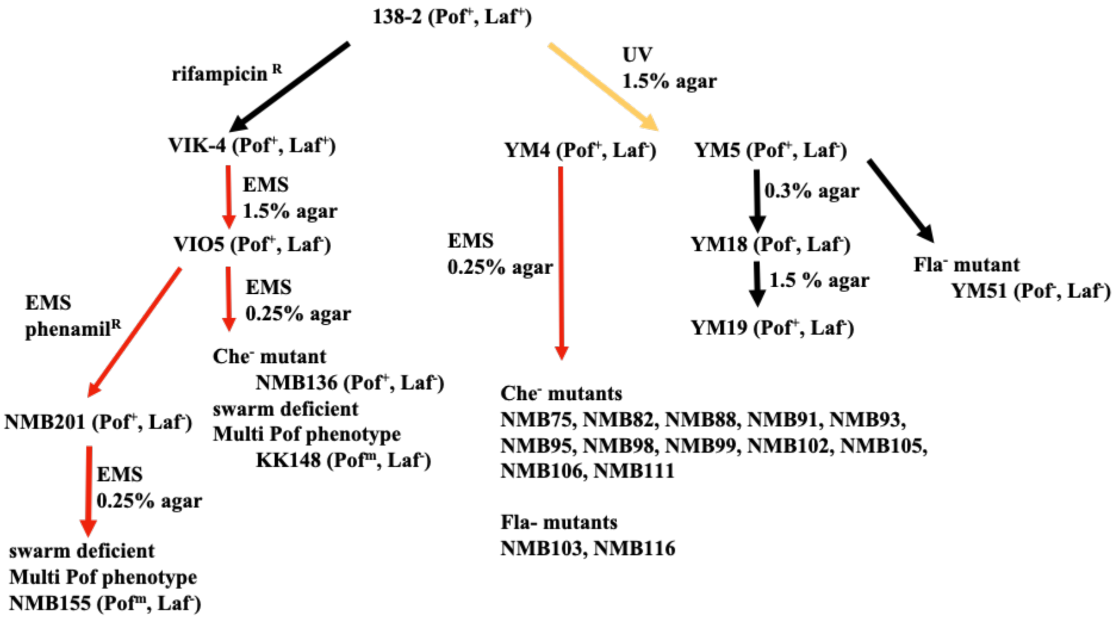


**Supplementary Fig.S1 The lineage of *V. alginolyticus* mutant strains**

*V. alginolyticus* strain 138-2 (Pof$^+$, Laf$^+$) was the parent for all the strains in this study. Strain VIO5 (Pof$^+$, Laf$^-$) was created by EMS mutagenesis of strain VIK-4, which is a spontaneous rifampicin-resistant strain obtained from strain 138-2. Strain NMB136 (Pof$^+$, Laf$^-$), NMB155 (Pof$^m$, Laf-) and strain KK148 (Pof$^m$, Laf-) were swarm-deficient mutants derived from strain VIO5. Strain YM4 (Pof$^+$, Laf$^-$) and YM5 (Pof$^+$, Laf$^-$) were obtained by UV-treatment of strain 138-2 and subsequent selection step on 1.5% agar plate. Strain YM4 was used for the flagella-deficient mutant selection using EMS mutagenesis. Strain YM18 (Pof$^-$, Laf$^-$) and revertant strain YM 19 (Pof$^+$, Laf$^-$) were derived from a low-concentration (0.3%) agar plate culture of strain YM5 and a subsequent normal-concentration (1.5%) agar plate culture of strain YM18. Strain YM51 (Pof$^-$, Laf$^-$) was derived from strain YM5. The red lines represent mutagenesis by EMS, the yellow line represents mutagenesis by UV irradiation and the black lines represent spontaneous mutations.

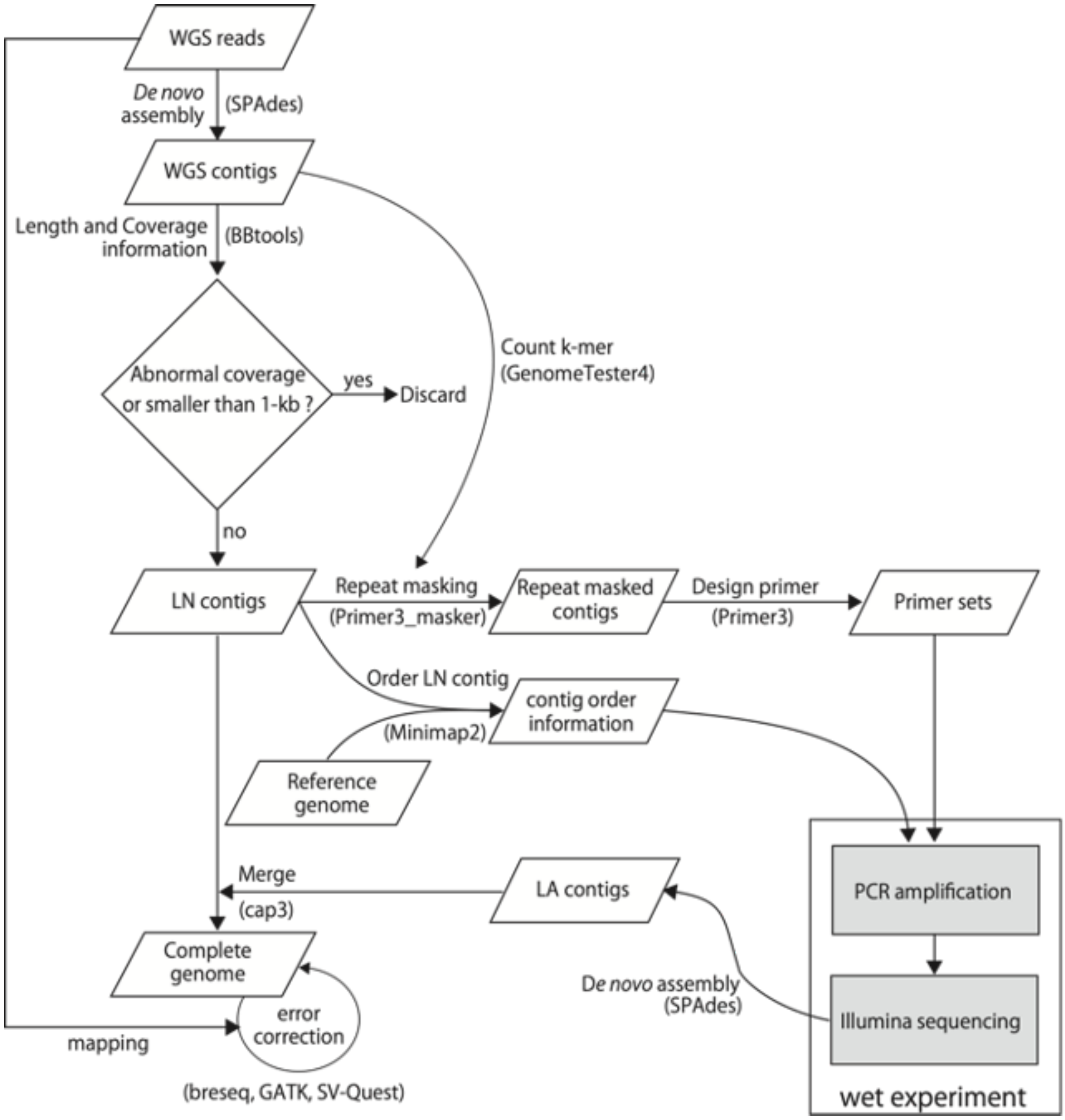


**Supplementary Fig.S2 Work-flow for complete genome sequencing using short read sequences**

Quality-filtered whole genome shotgun sequencing reads (WGS Reads) are used as input data for this workflow. WGS reads are de novo assembled and filtered based on length and average coverage. The remaining long and non-repetitive contigs are called LN contigs (Long and Normal coverage contigs). The LN contigs are mapped to a very closely related genome to determine the order and orientation of each contig. Fragments between LN contigs are amplified by PCR using primers designed at both ends of the LN contigs. Amplified fragments (2kbp~10kbp) are multiplex-sequenced with MiSeq sequencer. Paired-end sequencing reads in each sample were locally assembled. These LA (Locally Assembled contig of PCR fragment) contigs were connected with LN contigs to produce two closed genome structures. White and grey boxes represent computational experiments and wet experiments, respectively.

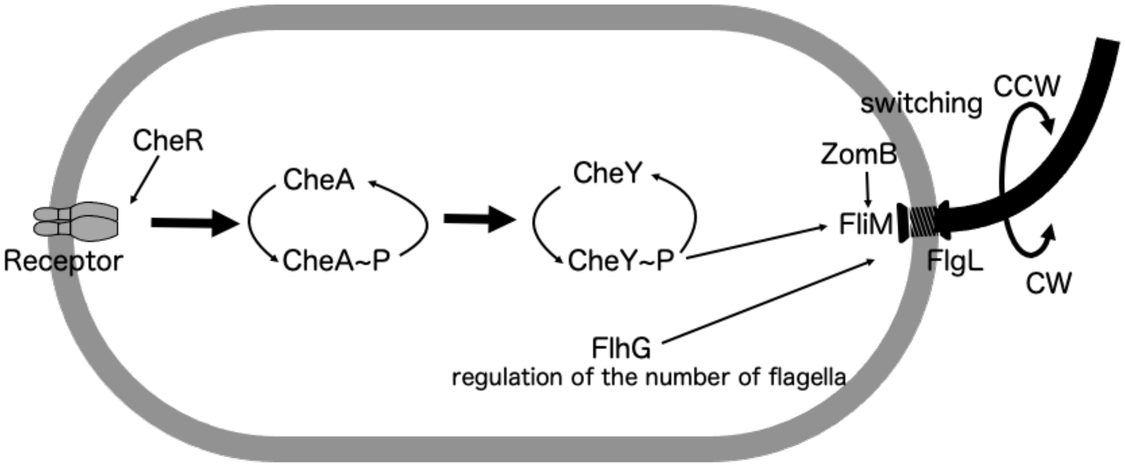


***cheR*:NMB75**

***cheA*:NMB82, 98**

***cheY*:NMB93, 136**

***fliM*:NMB88, 95, 99, 102, 106, 155**

***flhG*:KK148,NMB111**

***zomB*:NMB91,98**

***flgL*:NMB103,116**

**Supplementary Fig.S3 Gene mutations in the swarm-deficient strains found in this study**

Several components to control the flagellar rotation from chemoreceptors to flagella were schematically depicted. Seventeen mutant strains analyzed in this study were found to have genetic mutations within these components. The genes and the names of the mutant strains are listed at the bottom.
